# Agreement between Garmin and Ultrahuman wearables and their association with continuous glucose monitoring: an n-of-1 study

**DOI:** 10.64898/2026.10.13.758757

**Authors:** Mario Lovrić, Morten Arendt Rasmussen, Dejan Strbad

## Abstract

**Background:** Consumer wearables and continuous glucose monitors (CGM) are increasingly worn together outside clinical settings. Few studies have examined whether two wearables worn at the same time agree with each other, and whether either tracks glucose metrics derived from CGM.

**Objective:** In one adult under free-living conditions, we examined (1) how well sleep, heart rate, heart rate variability (HRV) and activity metrics from a Garmin smartwatch and an Ultrahuman ring agreed with each other, and (2) how these metrics related to glucose measured by CGM.

**Methods:** One healthy adult with no diagnosed diabetes wore a Garmin Fenix 8 watch on the wrist, an Ultrahuman Ring AIR and an Abbott FreeStyle Libre CGM at the same time for 12 days (29 July to 9 August 2026; 1101 glucose readings). We assessed agreement between the devices with Pearson correlation and Bland-Altman analysis (mean bias and 95% limits of agreement, LoA) for daily metrics and for heart rate within the day, and with Cohen’s kappa for sleep stages, minute by minute. We related the wearable metrics to mean glucose, glucose variability (coefficient of variation, CV%) and time in range (TIR, 3.9 to 10.0 mmol/L), reading by reading (over the whole record and separately for sleep and waking hours) and day by day. All correlations are descriptive. Because the Ultrahuman ring recorded no data for 22 hours on 3 August and its daytime heart rate behaved differently afterwards, heart-rate analyses are reported for the whole period and separately before and after that gap. The daily results were checked by leaving out one day at a time and by excluding the first 24 hours of sensor wear.

**Results:** The devices agreed closely on total sleep duration (r=0.94; bias −0.02 h, 95% LoA −0.56 to 0.51 h). Daily steps correlated strongly (r=0.99 over the 11 days with data from both devices), but Garmin counted more steps on every day (mean 11,157 vs. 8,929 steps per day; bias +2228 steps). Heart rate agreed moderately during sleep (r=0.68). While awake, agreement was moderate before the gap (r=0.54) and absent after it (r=0.08), when Ultrahuman’s daytime heart rate stayed near 70 beats per minute. The devices assigned the same sleep stage in 50.0% of 4525 minutes (kappa=0.28). Mean glucose was 5.13±0.82 mmol/L and TIR was 97.5%. Of all wearable signals, heart rate was most closely associated with glucose (Garmin r=0.43 over all readings), but the correlation fell to 0.34 within sleep and 0.21 within waking hours, and before the gap the two devices’ waking correlations were alike (Garmin 0.25, Ultrahuman 0.24). Across the 12 days, correlations between night-time wearable metrics and glucose variability weakened markedly when the first 24 hours of sensor wear were excluded (|r| fell from ≥0.59 to ≤0.34), and TIR was 100% on 8 of the 12 days, which left it little room to vary.

**Conclusions:** The two wearables agreed on how long the participant slept but not on how the night was staged, and their heart-rate agreement depended on whether the participant was asleep or awake and on the period of wear. Of all wearable signals, heart rate was most closely associated with glucose, mainly because both are lower at night and higher during the day. The associations across days rested largely on the first night of sensor wear. In this single participant, the devices behaved differently by device and by period of wear, so studies that combine different wearables, or relate them to CGM, should check for such differences before pooling data.

## Introduction

Consumer wearables such as smartwatches and smart rings estimate sleep, heart rate, heart rate variability (HRV) and physical activity from optical and motion sensors, and many people wear them day and night. Their sleep estimates have mostly been validated against polysomnography (PSG), the clinical reference method for sleep. These studies generally find that wearables detect sleep well but detect periods of wake during the night poorly, and that they classify the stages of sleep (light, deep and rapid eye movement [REM] sleep) inconsistently (Chinoy et al., 2021). In a recent comparison of six wrist-worn devices against PSG, all detected more than 90% of sleep epochs but only 29% to 52% of wake epochs, and Cohen’s kappa for epoch-by-epoch sleep staging ranged from 0.21 for the Garmin Vivosmart 4 to 0.53 for the Apple Watch Series 8 (Schyvens et al., 2025). Studies at home against electroencephalography and in the laboratory against PSG likewise found that most devices, including the Oura ring, estimate total sleep time better than the time spent in each stage, with the Garmin Vivosmart among the least accurate for total sleep time, and the laboratory study found that the error in stage durations often depended on how long the stage lasted (Kainec et al., 2024; Stone et al., 2020). Heart-rate metrics also differ between devices. Against an electrocardiography (ECG) chest strap, Oura rings measured nightly resting heart rate and HRV most accurately, while a Garmin watch measured HRV less accurately (Dial et al., 2025). Many devices combine these signals into readiness or recovery scores, such as Garmin Body Battery and Ultrahuman Dynamic Recovery, using formulas that the manufacturers do not disclose (Doherty et al., 2025). Most users, however, have no reference device and can only compare one wearable with another. The validity and reliability of consumer wearables have been studied mainly for steps, heart rate and energy expenditure, mostly in controlled laboratory settings (Fuller et al., 2020); how well two different consumer wearables worn at the same time agree in daily life, and whether their agreement changes over the day, has been studied less.

Continuous glucose monitors (CGM) were developed for diabetes care, but people without diabetes increasingly use them to see how diet, physical activity and stress affect their glucose (Holzer et al., 2022). For diabetes, an international consensus defines standard CGM metrics, including time in range (TIR), the share of readings between 3.9 and 10.0 mmol/L (Battelino et al., 2019). In people without diabetes, glucose stays between 3.9 and 7.8 mmol/L about 96% of the time, with a coefficient of variation (CV) of about 17% when measured with a Dexcom sensor (Shah et al., 2019), although some people with normal results on standard tests reach prediabetic levels after meals (Hall et al., 2018). In diabetes and related disorders of glucose regulation, wearable signals carry information about glucose: heart rate and HRV change during hypoglycemia (Lehmann et al., 2023), sleep measured by a Garmin watch differs on nights with nocturnal hypoglycemia (Alsuhaymi et al., 2026), and a heart-rate peak at meal onset improves CGM-based meal detection in people with post-bariatric hypoglycemia (Cossu et al., 2026). Few studies have related wearable sleep, recovery, HRV and activity data to CGM metrics in people without diabetes, in whom glucose varies little. In that setting, glucose and heart rate both follow the day/night cycle and both rise after meals, so a correlation between them can arise from shared timing alone.

We therefore followed one adult without diabetes who wore a Garmin Fenix 8 smartwatch, an Ultrahuman Ring AIR and a CGM at the same time for 12 days. We asked (1) how well the two wearables agreed on sleep, heart-rate, HRV and activity metrics (research question 1, RQ1), and (2) how these metrics related to glucose metrics derived from CGM (RQ2). Because the ring recorded no data for 22 hours midway through the period, we also asked whether the answers differed before and after that gap.

## Materials and Methods

### Study design and participant

This was an observational study of a single participant (n-of-1) under free-living conditions. The participant, one of the authors, was a healthy adult with no diagnosed diabetes. No intervention was applied: diet, sleep and exercise followed the participant’s usual routine.

### Devices, data export and matching

The participant wore an Abbott FreeStyle Libre sensor on the back of the upper arm, a Garmin Fenix 8 watch (Sensor Hub 40.08; BMX 25.09) on the wrist and an Ultrahuman Ring AIR (firmware R1.01.01.01) on a finger from 29 July to 9 August 2026 (12 calendar days and nights). The CGM record runs from 17:49 on 29 July to 07:20 on 9 August (253.5 hours), so the first and last calendar days are partial (38 and 29 readings). The sensor stores a glucose value (mmol/L) about every 15 minutes (historic readings, n=941) and gives an additional reading whenever it is scanned (scan readings, n=160); we used both. After removing records duplicated when the sensor was paired a second time and empty placeholder records, and keeping one reading per timestamp, 1101 glucose readings remained: a median of 104 per full calendar day (range 76 to 133), with four gaps of more than one hour (61 to 327 minutes, all between late afternoon and shortly after midnight). The first 24 hours of wear, when readings are least accurate, are examined separately in the sensitivity analyses. The participant logged food and coffee intake on an Apple iPhone with the Timelines app (https://timelines.app/).

From each wearable we exported the intraday records (heart rate, HRV, skin temperature, steps, and Garmin’s stress score and Body Battery) through the vendors’ application programming interfaces (API) with an in-house Python workflow, at each device’s own resolution: about every 6 minutes for Ultrahuman and every 2 minutes for Garmin. Each glucose reading was matched to the nearest wearable record within that interval, and readings without a match were left missing. Daily summaries and sleep-stage intervals came from each device’s own export. Both devices assign a night to the date on which the participant woke up. The participant remained in one time zone throughout. The monitoring period, glucose metrics and data completeness are summarized in Table 1.

**Table 1.** Monitoring overview.

| Metric | Value |
| --- | --- |
| Monitoring window | 29 Jul to 9 Aug 2026 (12 days/nights) |
| Glucose readings, n | 1101 |
| Mean glucose, mmol/L | $5.13 \pm 0.82$ |
| Glucose CV, % | 16.0 |
| Time in range (3.9 to 10.0 mmol/L), % | 97.5 |
| Time below range ($<3.9$ mmol/L), % | 2.5 |
| Time above range ($>10.0$ mmol/L), % | 0.0 |
| Nights with paired UH/Garmin data | 12/12 |
| Mean total sleep, UH / Garmin, h | $6.06 \pm 0.81$ / $6.04 \pm 0.82$ |
| Mean daily steps, UH / Garmin | 8192 / 10,618 |
| Workouts logged, n (total min) | 12 (281) |
| Food events logged, n | 39 |
| Coffee events logged, n | 16 |

### Glucose metrics and device indices

Glucose metrics were computed for each calendar day (00:00 to 24:00, local time) and for the whole period: mean glucose, coefficient of variation (CV%, 100 × standard deviation [SD] / mean) and TIR (percentage of readings from 3.9 to 10.0 mmol/L), over all readings, historic and scan, without time weighting. This range is the consensus target for adults with type 1 or type 2 diabetes; the consensus sets no target range for people without diabetes and recommends at least 14 days of data with at least 70% sensor wear (Battelino et al., 2019). All TIR values in this paper use this range; the 3.9 to 7.8 mmol/L range cited in the Introduction was not used. Resting heart rate, night HRV, sleep score, Body Battery and the daily Body Battery charged (the amount by which Garmin’s Body Battery index rose over a day, mostly during sleep) are calculated by each manufacturer, and the formulas behind the composite scores (sleep score and Body Battery) are not disclosed (Doherty et al., 2025).

### Agreement between the watch and the ring (RQ1)

We compared eight daily metrics (total sleep time; sleep score; minutes of deep, light and REM sleep; resting heart rate; night HRV; daily steps) and intraday heart rate at the glucose timestamps where both devices had a matched reading. Maximal oxygen uptake (VO2max) was not compared because Garmin reported it on one day only, and Ultrahuman’s daily step total for 3 August (84 steps, recorded while the ring was off for about 22 hours) was excluded, leaving 11 days for steps. For each metric, we calculated the Pearson correlation, the mean bias (Garmin minus Ultrahuman) with its 95% confidence interval (CI), and the 95% limits of agreement (LoA; bias ± 1.96 SD of the differences) as described by Bland and Altman (Bland and Altman, 1986). Pearson r describes association, not agreement; the bias and LoA describe agreement. With 12 days, each limit of agreement has a 95% CI of about ±1.1 SD of the differences, so the LoA are illustrative. Heart-rate pairs were classed as asleep when either device had scored a sleep stage at that time, including wake within a sleep period, and as awake otherwise. Waking pairs were further split at 4 August, when Ultrahuman data resumed after the gap. Sleep stages (awake, light, REM, deep) were compared minute by minute over the full stage records, as percentage agreement and Cohen’s kappa, interpreted with the conventional labels of Landis and Koch: slight (≤0.20), fair (0.21 to 0.40), moderate (0.41 to 0.60), substantial (0.61 to 0.80) or almost perfect (>0.80) (Landis and Koch, 1977). Kappa depends on how common each stage is, so the stage distribution is reported with it. As a secondary comparison, we also compared the stages at the glucose timestamps that fell within a scored stage on both devices, because these are the time points used when glucose is related to sleep stage.

### Wearable signals and glucose (RQ2)

Within the day, we correlated each glucose reading with the matched wearable reading (Pearson r), first over all readings and then separately for readings taken asleep and awake, because glucose and heart rate both differ between sleep and waking hours. Because Ultrahuman’s daytime heart rate behaved differently after the data gap (see Results), the heart-rate correlations were also computed separately for 29 July to 2 August and for 4 to 9 August. The number of readings differs between wearable streams because each stream has its own sampling interval and coverage. To describe the daily rhythm, we computed mean glucose and each device’s mean heart rate, with 95% CI, for each clock hour. Across days, we correlated each night’s wearable summary with mean glucose, CV% and TIR on the date the night ended (n=12; n=11 for Ultrahuman steps). Ultrahuman active minutes were not analysed because the export reported 0 minutes on 10 of the 12 days. Because both devices assign a night to the wake-up date, each date includes the hours after midnight of that night, so this analysis does not separate night-time from daytime glucose. For daily correlations with |r| ≥ 0.50, we report the range of r when each day was left out in turn (leave-one-day-out) and r after excluding the first 24 hours of sensor wear. Overnight glucose was plotted against hours since sleep onset as recorded by Ultrahuman, and, to check whether within-sleep correlations reflect the time of night, each stream was also correlated with hours since sleep onset.

To check whether the intraday correlations reflected meals or exercise rather than a more direct link with glucose, we repeated them without readings taken within 90 minutes of a logged food or coffee event, or during or within 30 minutes after a logged workout. Finally, for each food and coffee event with at least two glucose readings before and after it, we compared mean glucose and heart rate in the 30 minutes before the event with their peak in the 150 minutes after and correlated the size of the glucose rise with the size of each device’s heart-rate rise across events. Heart rate was taken at the glucose-matched timestamps, so an event without at least two matched heart-rate readings in both windows has no heart-rate response. Events were often logged close together, so many 150-minute windows contain another logged event or a workout; we report the correlations over all events and over the events whose window contained neither.

### Statistical analysis

Glucose and wearable readings from one person form autocorrelated series, and the daily values are 12 consecutive days from the same person, so all correlations are descriptive: we report no p-values, significance thresholds or corrections for multiple comparisons, and the numbers of readings overstate the independent information in the data. We describe |r| below 0.30 as weak, 0.30 to 0.69 as moderate and 0.70 or more as strong; these are conventional labels, not tests. Missing data were not imputed. In the figures, glucose is smoothed with a centered 1-hour rolling mean and heart rate with a centered 30-minute rolling mean, and lines are broken at gaps in the data; all statistics used unsmoothed values. Analyses were done in Python (pandas, scipy, scikit-learn, matplotlib) and R (dplyr); the scripts are available with the data.

### Ethics

This was a self-monitoring study with a single participant, one of the authors (M.L.), who wore the devices as part of his own routine self-tracking and consented to the analysis and publication of his own data. All devices were commercially available, already in personal use and used as intended by the manufacturers: two non-invasive wearables and a minimally invasive CGM sensor. No other individuals were involved and no intervention was applied. No application was made to an ethics committee, because the study consists only of the author’s own observational data from consumer devices.

## Results

### Twelve days of glucose, heart-rate and sleep data

Over the full 12-day record (Figure 1), 1101 glucose readings were recorded (941 historic and 160 scan readings; mean 5.13±0.82 mmol/L; CV 16.0%; TIR 97.5%; Table 1). Glucose rose after logged meals and peaked at 9.1 mmol/L 50 minutes after a sweet snack on the first evening; it never exceeded 10.0 mmol/L, so all time outside the TIR range (3.9 to 10.0 mmol/L) was time below range. Of the 28 readings below 3.9 mmol/L, 24 fell during the first night of sensor wear (Figure 1A). The heart-rate traces of the two devices followed each other closely at night but diverged on several days, most visibly from 4 August onward, after Ultrahuman recorded no data for about 22 hours on 3 August (Figure 1B). Both devices exported complete daily summaries for all 12 nights, with a mean total sleep of 6.06±0.81 h for Ultrahuman and 6.04±0.82 h for Garmin. The participant logged 12 workouts (281 min in total), 39 food events and 16 coffee events.

**Figure 1.**
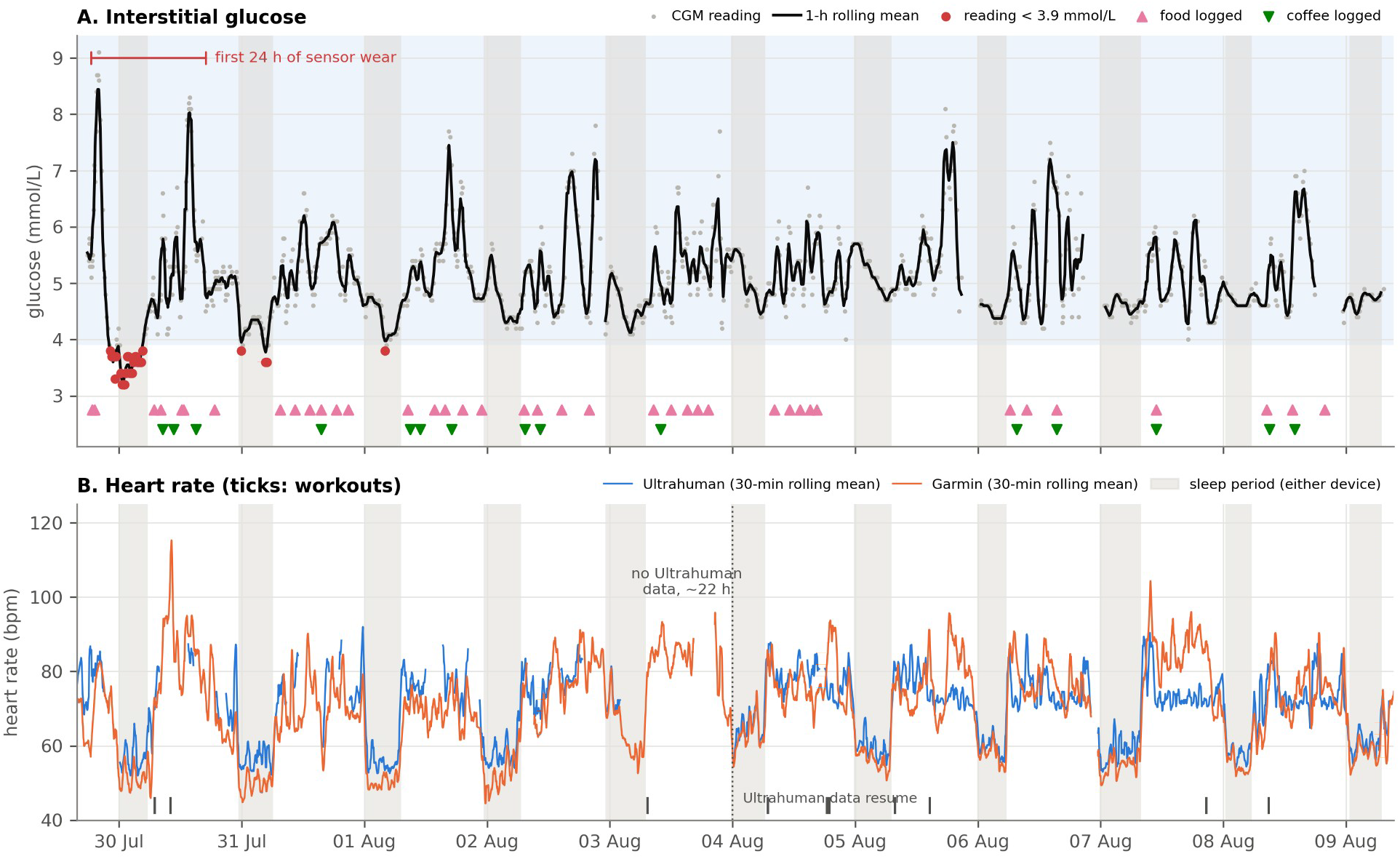
The 12-day record of glucose and heart rate. (A) Interstitial glucose: grey dots, individual CGM readings; black line, centered 1-hour rolling mean; red dots, readings below 3.9 mmol/L; blue band, 3.9 to 10.0 mmol/L range; triangles, logged food (pink) and coffee (green). The red bracket marks the first 24 hours of sensor wear. (B) Heart rate from Ultrahuman (blue) and Garmin (orange), centered 30-minute rolling means; ticks at the bottom mark logged workouts. Grey columns in both panels mark sleep periods (from the earlier bedtime to the later wake time of the two devices). Lines are broken at gaps in the data, including the ∼22-hour Ultrahuman gap on 3 August.

### Agreement between the watch and the ring (RQ1)

The Bland-Altman plots for the eight daily metrics reported by both devices are shown in Figure 2. They show which daily summaries could be used interchangeably (Table 2). Total sleep time agreed closely (r=0.945; bias −0.02 h, 95% CI −0.20 to 0.15 h; 95% LoA −0.56 to 0.51 h; Figure 2A). Daily steps correlated strongly (r=0.988 over the 11 days with data from both devices), but Garmin counted more steps on every day (mean 11,157 vs. 8,929 steps per day; bias +2228 steps, 95% CI 1574 to 2881, about 22% of the mean of the two devices; Figure 2H). On 3 August, the day the ring was off, Ultrahuman recorded 84 steps against Garmin’s 4699; with that day included, the bias would be +2427 steps. Sleep scores also correlated strongly (r=0.924), with Garmin 5 points lower on average (95% CI −9.5 to −0.9); because both scores are proprietary composites, this difference has no physiological meaning. Resting heart rate (r=0.566; Garmin 4.3 beats per minute [bpm] lower, 95% CI −5.9 to −2.8) and night HRV (r=0.620; Garmin 19 ms lower, 95% CI −25 to −13) correlated moderately. For night HRV, both limits of agreement lay below zero, and the difference shrank as mean HRV rose (Figure 2G). Light sleep (r=0.563; Garmin +39 min) and REM sleep (r=0.601; Garmin −37 min) correlated moderately, whereas deep-sleep minutes did not correlate at all (r=−0.155; 95% LoA −89 to 83 min, against a mean of about 80 min; Figure 2C). With 12 days, the limits of agreement themselves are imprecise: for total sleep time each limit has a 95% CI of ±0.30 h, and for deep sleep ±48 min.

**Table 2.** Garmin vs. Ultrahuman device agreement, daily and intraday metrics. Mean bias is given with its 95% CI. Daily steps exclude 3 August, when the ring was off for about 22 hours. UH, Ultrahuman.

| Metric | n | Pearson $r$ | Mean bias (Garmin – UH), 95% CI | 95% limits of agreement |
| --- | --- | --- | --- | --- |
| Total sleep (h) | 12 | 0.945 | –0.02 (–0.20 to 0.15) | –0.56 to 0.51 |
| Sleep score | 12 | 0.924 | –5.17 (–9.47 to –0.86) | –18.46 to 8.12 |
| Deep sleep (min) | 12 | –0.155 | –2.83 (–30.65 to 24.99) | –88.65 to 82.98 |
| Light sleep (min) | 12 | 0.563 | +38.75 (14.02 to 63.48) | –37.53 to 115.03 |
| REM sleep (min) | 12 | 0.601 | –37.42 (–61.80 to –13.03) | –112.64 to 37.81 |
| Resting HR (bpm) | 12 | 0.566 | –4.33 (–5.87 to –2.79) | –9.09 to 0.42 |
| Night HRV (ms) | 12 | 0.620 | –18.83 (–24.85 to –12.81) | –37.41 to –0.26 |
| Daily steps | 11 | 0.988 | +2227.5 (1574.2 to 2880.9) | 321.4 to 4133.7 |
| Intraday HR (bpm), all | 647 | 0.661 | –1.51 (–2.31 to –0.70) | –21.93 to 18.92 |
| Intraday HR, asleep | 200 | 0.676 | –4.49 (–5.30 to –3.68) | –15.94 to 6.96 |
| Intraday HR, awake | 447 | 0.270 | –0.17 (–1.26 to 0.92) | –23.04 to 22.70 |
| Intraday HR, awake, 29 Jul to 2 Aug | 180 | 0.539 | –4.58 (–6.01 to –3.15) | –23.63 to 14.47 |
| Intraday HR, awake, 4 to 9 Aug | 267 | 0.079 | +2.81 (1.37 to 4.24) | –20.61 to 26.23 |

**Figure 2.**
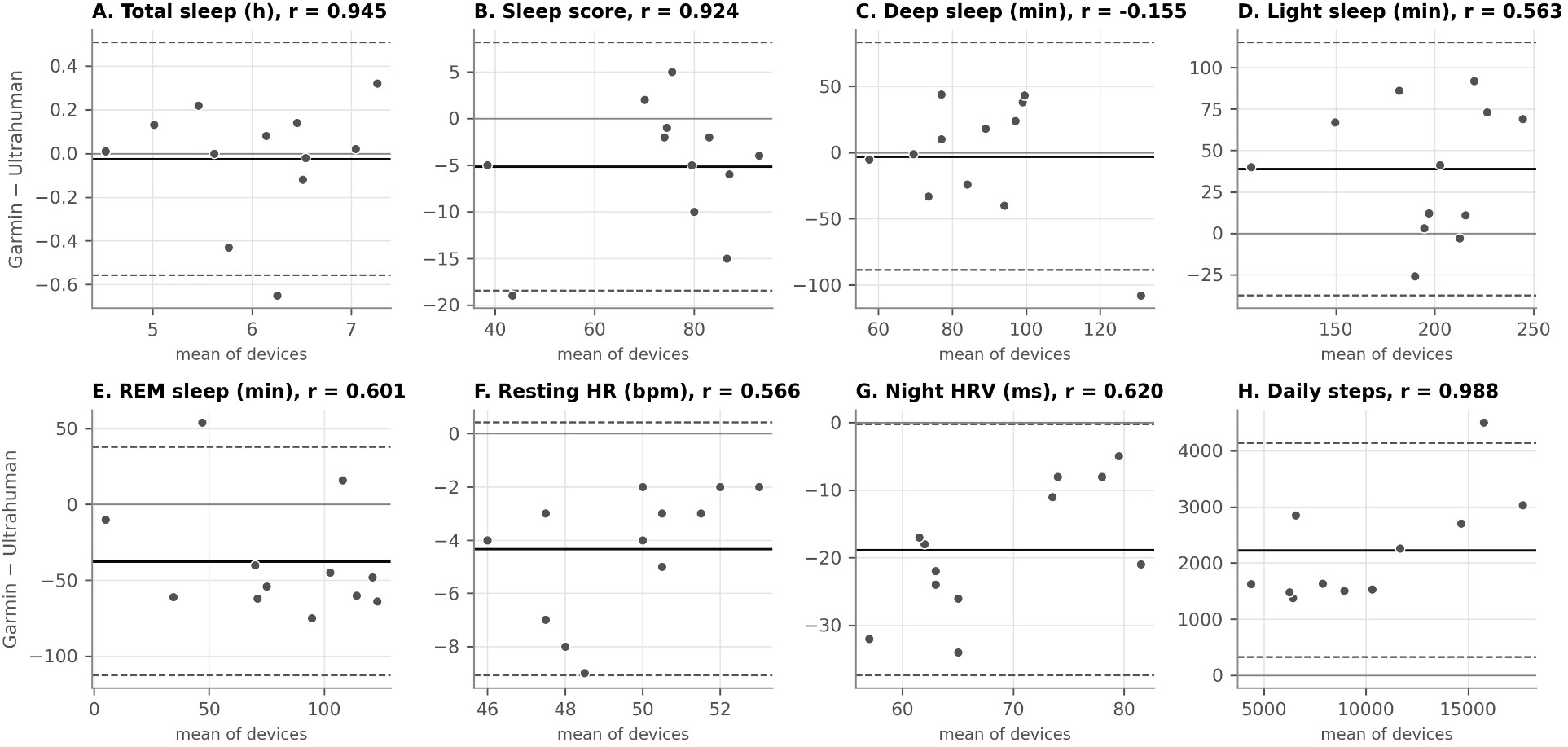
Agreement between Garmin and Ultrahuman daily metrics (Bland-Altman, n=12 days; 11 for daily steps, because 3 August was excluded). Each panel plots the Garmin − Ultrahuman difference against the mean of the two devices for one daily metric, with the Pearson r in the panel title. Solid black line, mean bias; dashed lines, 95% limits of agreement; grey line, zero difference. VO2max is not shown because Garmin reported it on one day only.

Beyond the daily summaries, heart-rate agreement appeared to change with time of day and over the study period (Figure 1B), so we broke down the 647 matched heart-rate readings by sleep or wake, clock time and date (Figure 3). During sleep (n=200), the two devices’ heart rates correlated moderately (r=0.676) and Garmin read 4.5 bpm lower (95% CI −5.3 to −3.7) with little scatter (SD of the differences 5.8 bpm). While the participant was awake, agreement depended on the period. Before the gap (29 July to 2 August, n=180), waking heart rates correlated moderately (r=0.539), with Garmin 4.6 bpm lower (95% CI −6.0 to −3.2; 95% LoA −23.6 to 14.5 bpm). After it (4 to 9 August, n=267) they no longer correlated (r=0.079), Garmin read 2.8 bpm higher (95% CI 1.4 to 4.2), and the 6-hour rolling median difference rose from around zero to about +15 bpm on 7 August (Figure 3D). Pooled over the whole period, waking agreement was weak (n=447; r=0.270; SD of the differences 11.7 bpm), and 45 of the 48 pairs outside the limits of agreement were waking pairs (Figure 3A and B; Table 2). The cluster of pairs in which Ultrahuman read near 70 bpm while Garmin read 70 to 100 bpm (Figure 3A) consisted of waking readings spread from morning to evening (Figure 3C), and 24 of the 25 pairs with Garmin above the upper limit came from 4 August onward (Figure 3D). From that date, 49% of Ultrahuman’s daytime (09:00 to 21:00) heart-rate samples read 69 to 73 bpm, compared with 8% before, and its heart rate no longer followed its own step count (r=0.02, compared with 0.39 before). At matched waking time points, Garmin heart rate correlated with Ultrahuman’s step count to the same degree in both periods (r=0.37), whereas Ultrahuman’s own heart rate did so before 4 August (r=0.42) but not after (r=0.06). The two readings in each pair were at most 3.7 min apart, and outlying pairs were no further apart than the rest, so differences in timing do not explain the disagreement. Over all 647 pairs the correlation was moderate (r=0.661), with Garmin 1.5 bpm lower on average (95% LoA −21.9 to 18.9 bpm), but this pooled estimate mixes a period of normal operation with a period in which the ring’s daytime heart rate was evidently not being measured correctly.

**Figure 3.**
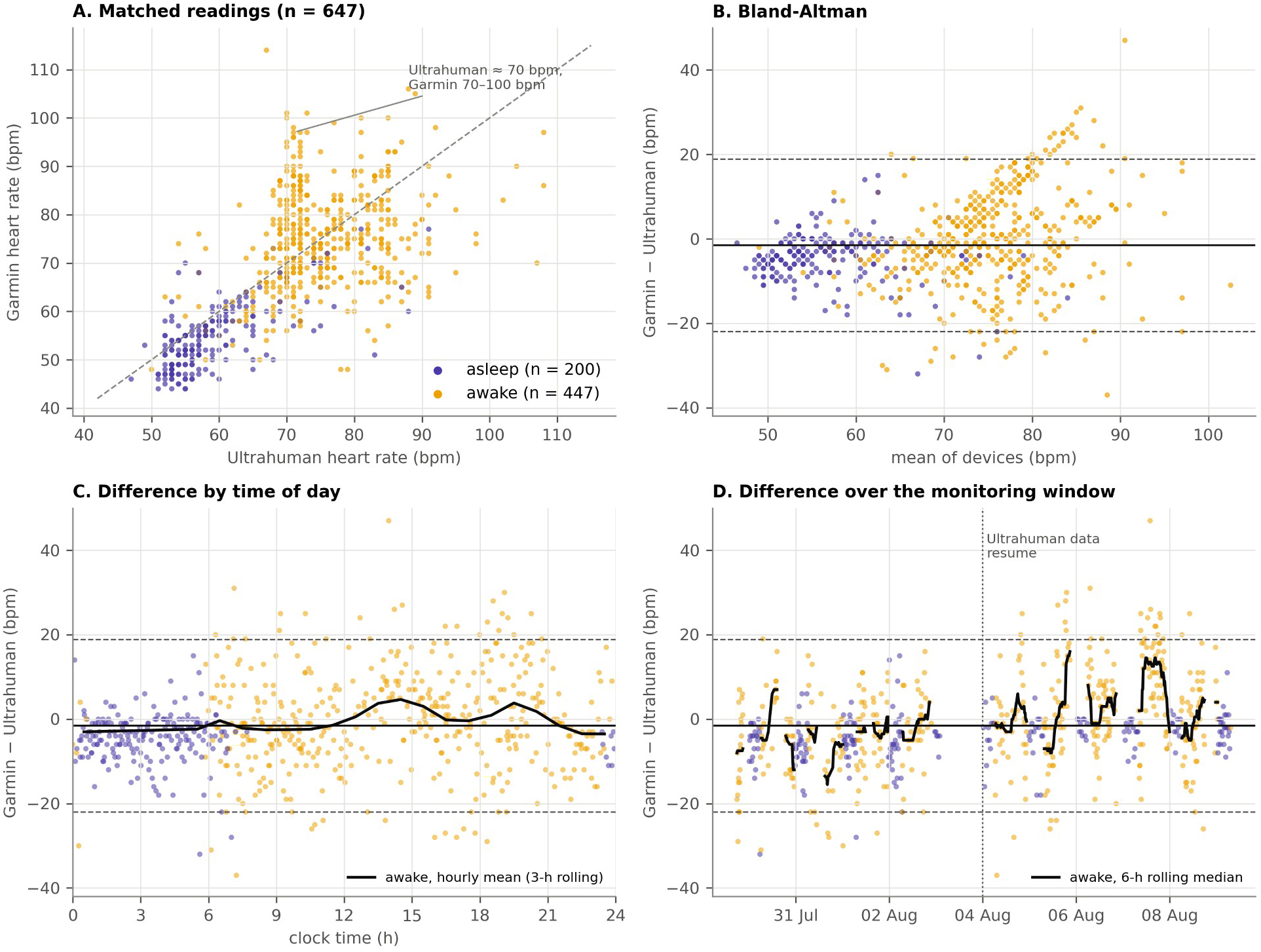
Intraday heart-rate agreement between Garmin and Ultrahuman by sleep/wake state, time of day and date (n=647 matched readings). (A) Garmin vs. Ultrahuman heart rate with the identity line; the annotation marks the cluster of waking readings with Ultrahuman near 70 bpm. (B) Bland-Altman plot. (C) Difference by clock time; black line, hourly mean of waking differences smoothed over 3 hours. (D) Difference over the monitoring window; black line, 6-hour rolling median of waking differences; dotted line, 4 August, when Ultrahuman data resumed after a ∼22-hour gap. Violet, asleep (either device assigned a sleep stage); yellow, awake. In B to D, solid line, mean bias; dashed lines, 95% limits of agreement.

Total sleep time agreed closely between the devices, but the sleep stages within each night did not. A confusion matrix shows where the two devices’ stage labels differed, and per-night hypnograms show whether the disagreement was spread across nights or concentrated in a few (Figure 4). Minute by minute over the 4525 minutes scored by both devices, they agreed on 50.0% (Cohen’s kappa=0.275, fair). By Ultrahuman’s scoring, light sleep made up 37% of these minutes, REM sleep 28%, deep sleep 26% and wake 9%, so kappa is dominated by light sleep. Most disagreement involved light sleep: of the minutes Ultrahuman scored as REM, Garmin scored 64% as light sleep, and of those Ultrahuman scored as deep, 33% (Figure 4A). Taking Ultrahuman’s stage as the reference, agreement was highest for light sleep (65%), followed by deep sleep (59%), wake (45%) and REM sleep (24%). Agreement varied between nights from 21% (2 August) to 67% (5 August) (Figure 4B). At the 272 glucose timestamps with a stage from both devices, the time points used later to relate glucose to sleep stage, agreement was similar (54.0%; kappa=0.331); only 26 of these time points were scored as wake by Ultrahuman. The hypnograms show that the devices broadly agreed on sleep onset, wake time and the block of deep sleep early in the night and disagreed mostly on the alternation of light and REM sleep in the second half of the night.

**Figure 4.**
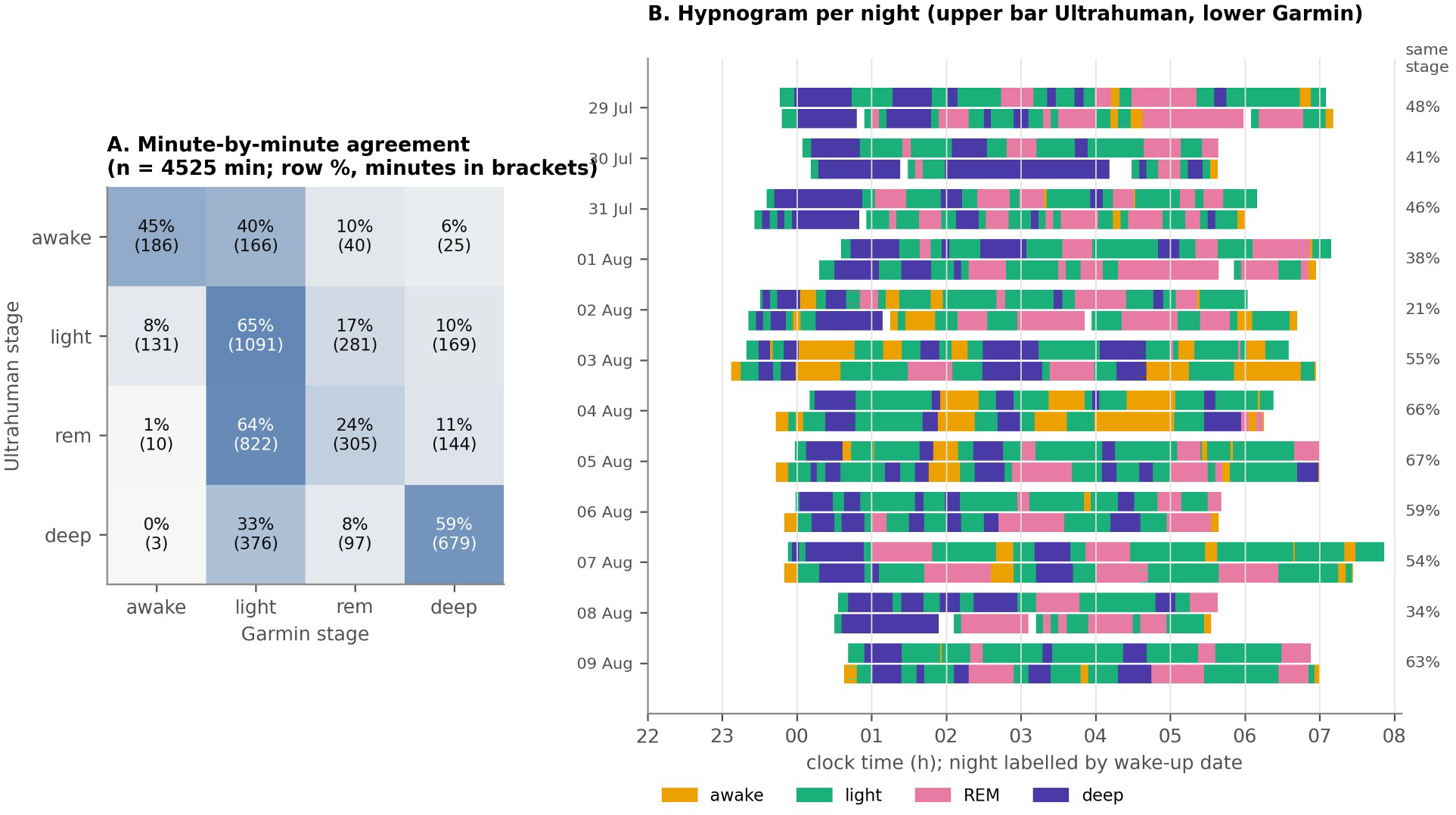
Sleep-stage agreement between Ultrahuman and Garmin. (A) Confusion matrix over the 4525 minutes scored by both devices; cells show row percentages (share of each Ultrahuman stage) with minutes in brackets. (B) Hypnograms for each of the 12 nights, labelled by wake-up date; for each night the upper bar is Ultrahuman and the lower bar Garmin, coloured by stage. The percentage on the right is the minute-by-minute agreement for that night.

### Wearable signals and glucose (RQ2)

#### Heart rate and glucose over the day

Glucose and heart rate share a day/night rhythm, so we plotted both by clock hour and compared correlations over all readings with correlations within sleep and within waking hours (Figure 5, Table 3). Mean glucose was lowest around 04:00 (4.40 mmol/L) and highest in the early afternoon (5.78 mmol/L at 14:00). Heart rate rose from about 55 to 59 bpm at night to about 76 to 77 bpm during the day on both devices (Figure 5A and B). Over all readings, Garmin heart rate had the strongest association with glucose (r=0.427, n=874), followed by Ultrahuman heart rate (r=0.377) and Garmin stress score (r=0.371). Within sleep or within waking hours, every heart-rate and stress correlation was weaker: Garmin heart rate, r=0.337 asleep and 0.208 awake; Ultrahuman heart rate, 0.257 and 0.090; Garmin stress, 0.305 and 0.135 (Figure 5C). For Ultrahuman skin temperature, the sign reversed, from r=−0.105 over all readings to 0.269 asleep and 0.171 awake. Garmin Body Battery was more strongly associated with glucose during sleep (r=−0.516) than over all readings (r=−0.138), and Ultrahuman HRV and steps were only weakly associated with glucose in every state. Before the Ultrahuman data gap, the two devices’ heart rates were associated with glucose to a similar degree (29 July to 2 August: Garmin r=0.483 over all readings, 0.208 asleep and 0.252 awake; Ultrahuman r=0.513, 0.262 and 0.243). After it, both weakened, and Ultrahuman’s waking correlation turned slightly negative (4 to 9 August: Garmin r=0.317, 0.216 and 0.157; Ultrahuman r=0.202, 0.190 and −0.091; Table 3). Ultrahuman’s near-zero waking correlation over the whole period therefore comes from the period after the gap.

**Table 3.** Intraday correlations between wearable streams and matched glucose, pooled and within sleep/wake state. Heart-rate rows are also given for the periods before (29 Jul to 2 Aug) and after (4 to 9 Aug) the Ultrahuman data gap.

| Metric | $r$, pooled (n) | $r$, asleep (n) | $r$, awake (n) |
| --- | --- | --- | --- |
| Garmin heart rate | 0.427 (874) | 0.337 (212) | 0.208 (662) |
| Ultrahuman heart rate | 0.377 (793) | 0.257 (266) | 0.090 (527) |
| Garmin heart rate, 29 Jul to 2 Aug | 0.483 (495) | 0.208 (123) | 0.252 (372) |
| Garmin heart rate, 4 to 9 Aug | 0.317 (379) | 0.216 (89) | 0.157 (290) |
| Ultrahuman heart rate, 29 Jul to 2 Aug | 0.513 (339) | 0.262 (116) | 0.243 (223) |
| Ultrahuman heart rate, 4 to 9 Aug | 0.202 (454) | 0.190 (150) | -0.091 (304) |
| Garmin stress | 0.371 (391) | 0.305 (137) | 0.135 (254) |
| Ultrahuman HRV | -0.162 (793) | -0.142 (268) | -0.150 (525) |
| Garmin body battery | -0.138 (529) | -0.516 (138) | -0.113 (391) |
| Ultrahuman skin temperature | -0.105 (968) | 0.269 (269) | 0.171 (699) |
| Ultrahuman steps | 0.055 (991) | 0.071 (269) | -0.054 (722) |

**Figure 5.**
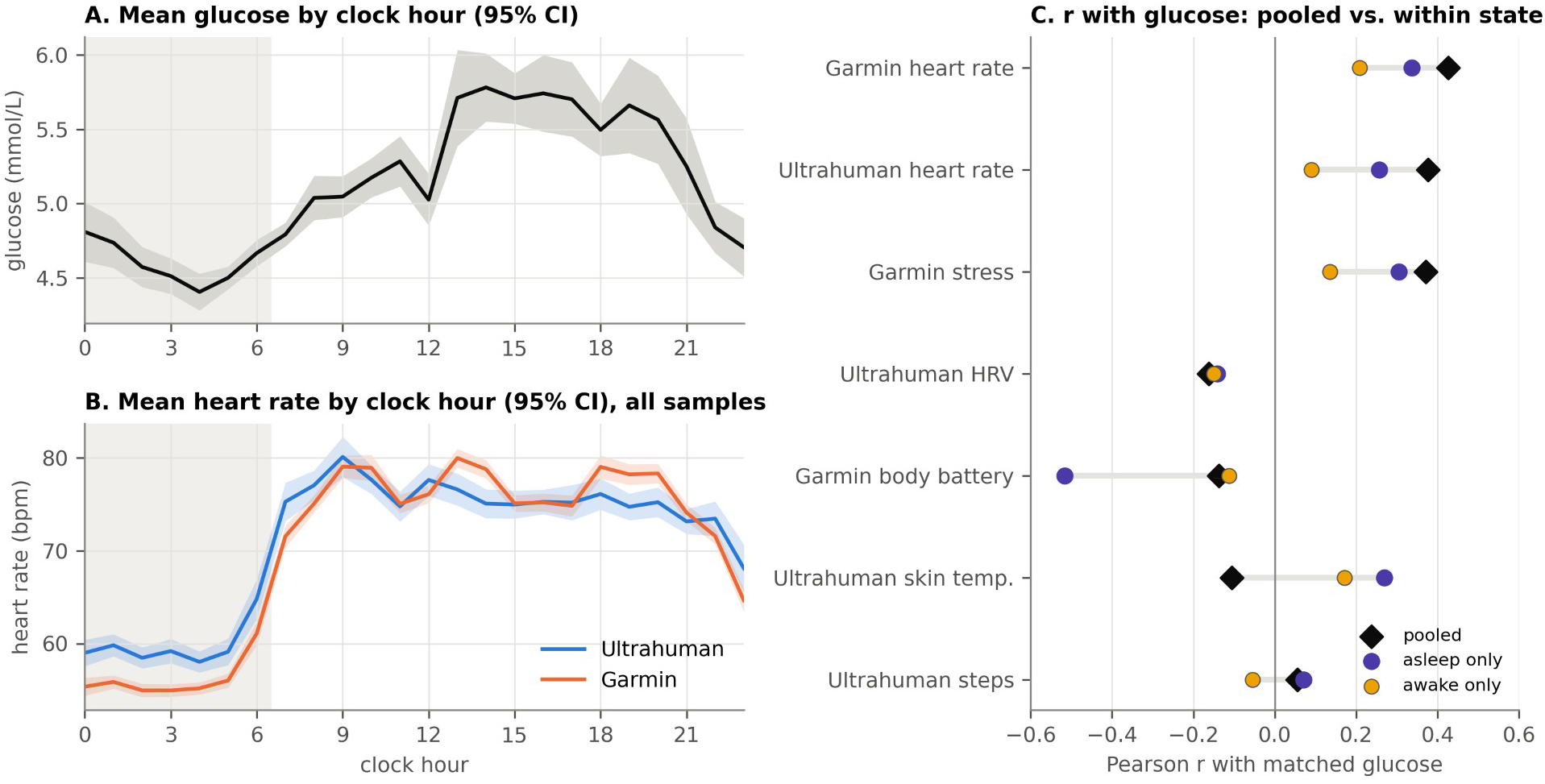
Shared day/night rhythm of glucose and heart rate, and its effect on their correlation. (A) Mean glucose by clock hour with 95% CI. (B) Mean heart rate by clock hour for Ultrahuman (blue) and Garmin (orange), all samples, with 95% CI. Grey shading in A and B marks the usual sleep hours. (C) Pearson r between matched glucose and each intraday wearable stream, pooled over the whole window (black diamond) and separately for readings taken asleep (violet) and awake (yellow); the grey bar spans the three estimates. Correlations are descriptive (autocorrelated series).

Meals and exercise raise both glucose and heart rate, which could explain the remaining correlations. Because food, coffee and workouts were logged (Table 1), we could test whether the heart-rate correlations depended on these events rather than on glucose itself. Readings within 90 minutes of a logged meal or coffee, or during or shortly after a workout, made up 33.3% of all matched readings and 44.8% of waking readings. Excluding them did not weaken the Garmin correlations (Table 4). The Garmin correlation became stronger, both over all readings (r=0.427 to 0.495) and while awake (r=0.208 to 0.269), whereas Ultrahuman’s already weak waking correlation fell further, to near zero (r=0.090 to 0.024). Garmin stress and Body Battery changed slightly in the same direction (stress while awake, r=0.135 to 0.152; Body Battery over all readings, r=−0.138 to −0.229). Meals and exercise are therefore unlikely to be the main driver of the Garmin heart-rate correlation with glucose. For Ultrahuman, the period split above suggests that the weak waking correlation reflects the ring’s behaviour after the gap rather than a difference in what it measures.

**Table 4.**
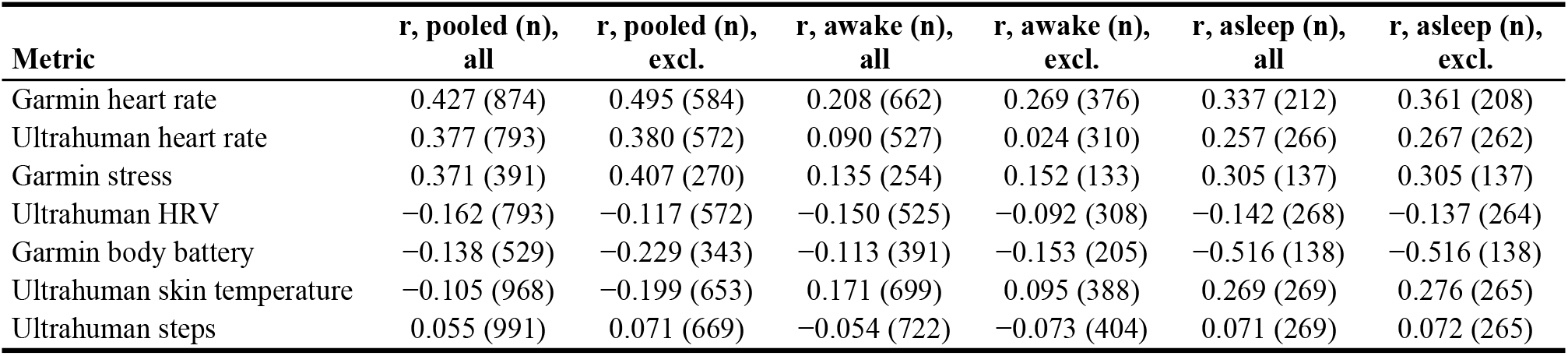
Intraday wearable-glucose correlations before and after excluding meal/coffee- and workout-proximate readings. Excluded: readings within 90 min of a logged food or coffee event, or during/within 30 min after a logged workout (33.3% of all matched readings, 44.8% of waking readings).

| Metric | r, pooled (n),<br>all | r, pooled (n),<br>excl. | r, awake (n),<br>all | r, awake (n),<br>excl. | r, asleep (n),<br>all | r, asleep (n),<br>excl. |
| --- | --- | --- | --- | --- | --- | --- |
| Garmin heart rate | 0.427 (874) | 0.495 (584) | 0.208 (662) | 0.269 (376) | 0.337 (212) | 0.361 (208) |
| Ultrahuman heart rate | 0.377 (793) | 0.380 (572) | 0.090 (527) | 0.024 (310) | 0.257 (266) | 0.267 (262) |
| Garmin stress | 0.371 (391) | 0.407 (270) | 0.135 (254) | 0.152 (133) | 0.305 (137) | 0.305 (137) |
| Ultrahuman HRV | -0.162 (793) | -0.117 (572) | -0.150 (525) | -0.092 (308) | -0.142 (268) | -0.137 (264) |
| Garmin body battery | -0.138 (529) | -0.229 (343) | -0.113 (391) | -0.153 (205) | -0.516 (138) | -0.516 (138) |
| Ultrahuman skin temperature | -0.105 (968) | -0.199 (653) | 0.171 (699) | 0.095 (388) | 0.269 (269) | 0.276 (265) |
| Ultrahuman steps | 0.055 (991) | 0.071 (669) | -0.054 (722) | -0.073 (404) | 0.071 (269) | 0.072 (265) |

Excluding readings around meals removes their effect but does not show how glucose and heart rate respond to each meal. We therefore also examined each logged food and coffee event separately, comparing the mean over the 30 minutes before the event with the peak over the 150 minutes after, for glucose and for each device’s heart rate (Table 5). Of the 54 events with enough glucose readings, 37 had a Garmin and 30 an Ultrahuman heart-rate response; the others lacked matched heart-rate readings in one of the windows. Food raised glucose by a median of 1.27 mmol/L (interquartile range [IQR] 0.48 to 2.09, n=38) and Garmin heart rate by 14.0 bpm (IQR 7.8 to 23.3, n=23). Coffee raised glucose less (median 0.40 mmol/L, IQR 0.05 to 0.73, n=16) but heart rate by a similar amount (9.5 bpm, IQR 5.0 to 20.8, n=14). Across events, a larger glucose rise went with a larger Garmin heart-rate rise, moderately for food (r=0.554, n=23) and strongly for coffee (r=0.848, n=14; all events together, r=0.623, n=37), and similarly before and after the Ultrahuman data gap (r=0.596, n=24, and r=0.659, n=13). Removing any single event changed these correlations by no more than 0.09. Ultrahuman’s heart-rate rise showed no such relationship in either period (all events, r=−0.058, n=30; before the gap, r=−0.040, n=15). These correlations should be read with caution: 32 of the 54 events had another logged event within the 150-minute window and 5 had a workout, so neighbouring responses overlap, and a peak compared with a 30-minute mean is biased upward in a noisy series. Among the 21 events whose window contained neither another event nor a workout, the Garmin correlation was weak (r=0.198, n=12; Ultrahuman, r=−0.033, n=12).

**Table 5.** Meal-locked glucose and heart-rate response to logged food and coffee events. Baseline, mean over the 30 min before the event; rise, post-event peak (within 150 min after) minus baseline. Correlation columns relate each event’s glucose rise to its heart-rate rise. Heart rate is taken at glucose-matched timestamps; events without at least two matched heart-rate readings in both windows have no heart-rate response, hence the smaller n.

| Metric | Food, median [IQR]<br>(n) | Coffee, median [IQR]<br>(n) | r with glucose<br>rise, all events (n) | food only<br>(n) | coffee only<br>(n) |
| --- | --- | --- | --- | --- | --- |
| Glucose rise (mmol/L) | 1.27 [0.48 to 2.09] (38) | 0.40 [0.05 to 0.73] (16) | — | — | — |
| Garmin HR rise (bpm) | 14.0 [7.8 to 23.3] (23) | 9.5 [5.0 to 20.8] (14) | 0.623 (37) | 0.554 (23) | 0.848 (14) |
| Ultrahuman HR rise (bpm) | 10.9 [2.8 to 14.7] (20) | 11.9 [9.8 to 16.8] (10) | -0.058 (30) | 0.031 (20) | -0.346 (10) |

#### Wearable summaries and glucose from day to day

Across days, we related each night’s wearable summary to glucose on the same date. To judge the few large correlations against the rest, all 39 are shown together in Figure 6. Ultrahuman night resting heart rate had the largest: r=0.89 with TIR and r=−0.81 with CV%, meaning that nights with a lower resting heart rate went with more variable glucose and less time in range on the same date. The correlations for Garmin resting heart rate were smaller (r=0.44 with TIR, r=−0.59 with CV%). Garmin Body Battery charged and night HRV were moderately associated in the same direction (|r| 0.61 to 0.67), Ultrahuman daytime HRV had |r| ≤ 0.32, and sleep duration and sleep score had |r| ≤ 0.40 on either device. Days with more steps on either device had lower mean glucose (Ultrahuman, r=−0.66 over 11 days; Garmin, r=−0.56).

**Figure 6.**
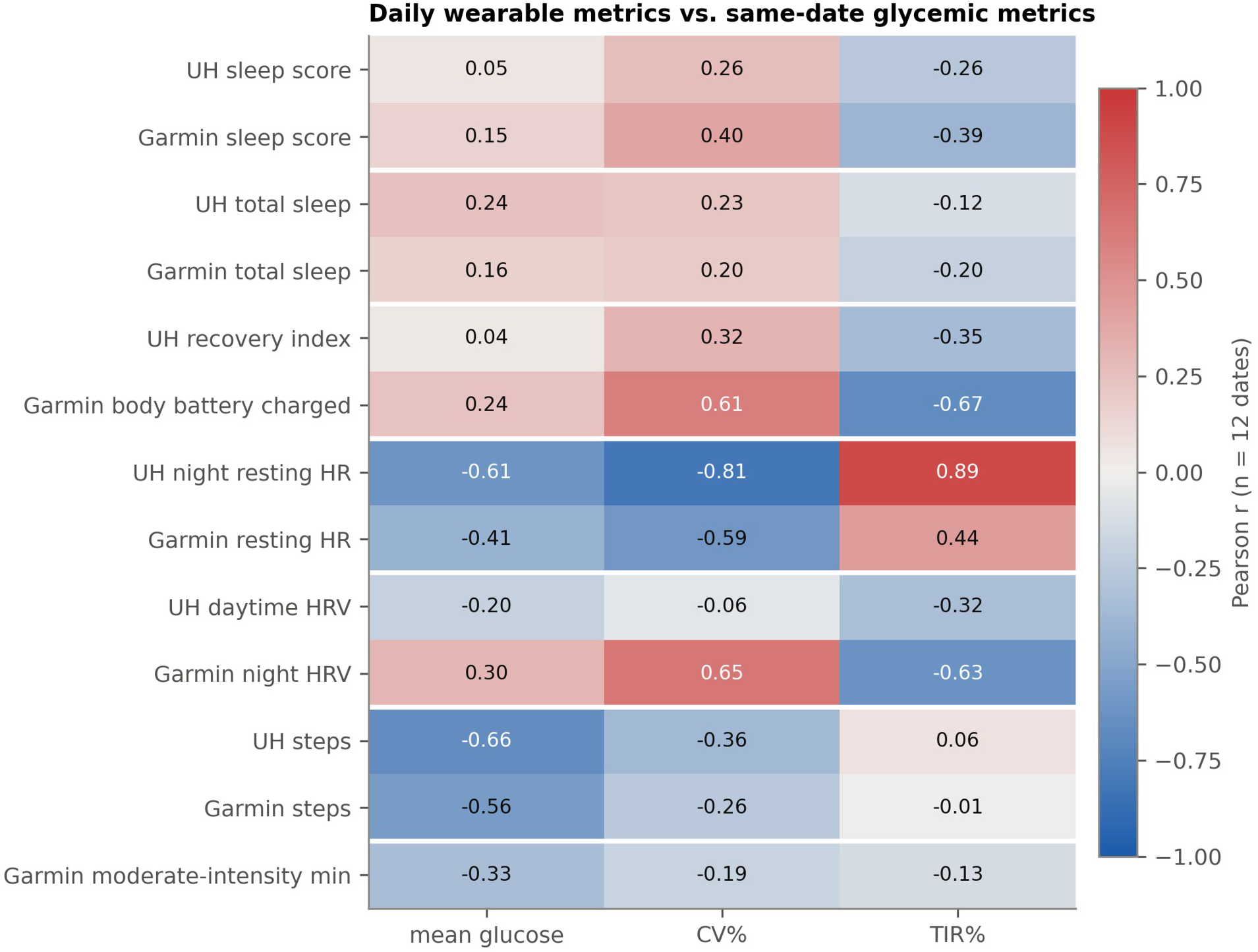
Daily wearable metrics vs. same-date glycemic metrics. Pearson r between each daily wearable metric (rows, grouped in device pairs) and glucose mean, CV% and TIR for the calendar date on which the night ended (n=12 dates; 11 for Ultrahuman steps). Blue, negative; red, positive. All 39 coefficients are descriptive: the dates are consecutive days from one person, so no significance threshold applies. UH, Ultrahuman; HR, heart rate.

With only 12 dates, a single unusual date can drive such correlations. Daily TIR was 100% on 8 of the 12 dates, and the two lowest values fell on 29 and 30 July (84% and 86%), the dates spanning the first night of sensor wear; TIR therefore varied on only four dates, and any correlation with it rests on those. We tested how much each daily association depends on individual dates (Figure 7, Table 6). Leaving out one date at a time changed most correlations by less than 0.2, but excluding the first 24 hours of sensor wear markedly weakened every correlation with CV% (|r| fell from ≥0.59 to ≤0.34; Ultrahuman night resting heart rate, r=−0.81 to 0.08; Garmin resting heart rate, r=−0.59 to −0.34; Figure 7A). The TIR correlations changed less (Ultrahuman night resting heart rate, r=0.82 after the exclusion), but after this exclusion TIR fell below 100% on only three dates, and by no more than 2.7 percentage points, so these coefficients describe two or three dates rather than a relationship across days. The scatterplots show that the two dates of the first sensor night combined the lowest resting heart rate with the lowest TIR and the highest CV%, and so account for most of each correlation (Figure 7B and C).

**Table 6.** Daily associations between wearable and glycemic metrics with |r| ≥ 0.50 (n=12 nights; 11 for Ultrahuman steps), with sensitivity analyses. LODO, range of r across leave-one-day-out fits (n=11 each); excl. first 24 h, r after excluding glucose readings from the first 24 hours of sensor wear (n=11 dates).

| Metric | Glycemic target | Pearson $r$ | LODO range | Excl. first 24 h |
| --- | --- | --- | --- | --- |
| Ultrahuman night resting HR | TIR | 0.89 | 0.79 to 0.92 | 0.82 |
| Ultrahuman night resting HR | CV% | -0.81 | -0.87 to -0.64 | 0.08 |
| Garmin body battery charged | TIR | -0.67 | -0.77 to -0.56 | -0.78 |
| Garmin HRV (night avg) | CV% | 0.65 | 0.51 to 0.71 | 0.13 |
| Garmin HRV (night avg) | TIR | -0.63 | -0.75 to -0.47 | -0.66 |
| Garmin body battery charged | CV% | 0.61 | 0.46 to 0.67 | 0.00 |
| Ultrahuman night resting HR | mean glucose | -0.61 | -0.80 to -0.27 | -0.18 |
| Garmin resting HR | CV% | -0.59 | -0.67 to -0.43 | -0.34 |
| Garmin daily steps | mean glucose | -0.56 | -0.66 to -0.47 | -0.50 |
| Ultrahuman daily steps | mean glucose | -0.66 | -0.77 to -0.55 | -0.58 |

**Figure 7.**
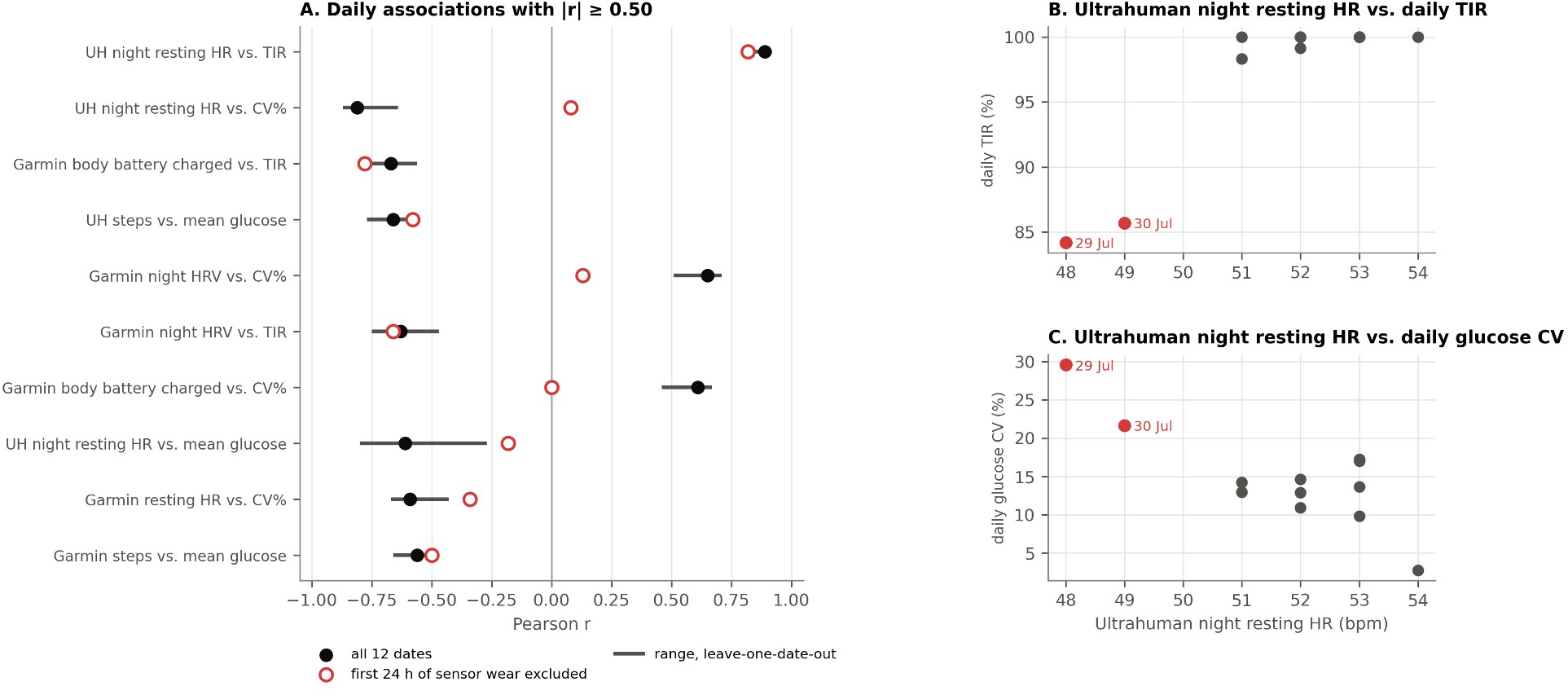
Robustness of the daily associations. (A) For each daily association with |r| ≥ 0.50: black dot, r over all 12 dates (11 for Ultrahuman steps); grey bar, range of r when each date is left out in turn; open red circle, r after excluding the first 24 hours of sensor wear. (B, C) Ultrahuman night resting heart rate against daily TIR (B) and daily glucose CV% (C); red points are 29 and 30 July, the dates that include the first night of sensor wear.

#### Glucose by sleep stage and time of night

The daily analysis mixes night-time and daytime glucose, so we also examined glucose during sleep directly, asking whether differences between sleep stages reflect the stages themselves or the time of night at which they occur (Figure 8). Median glucose was 4.9 mmol/L at time points scored as awake, 4.6 mmol/L in light sleep, 4.4 mmol/L in REM sleep and 4.6 mmol/L in deep sleep. These medians were identical for both devices’ staging, even though the devices often disagreed on the stage itself. Deep sleep had the widest spread and the lowest values, all from the first sensor night (Figure 8A). Excluding that night, overnight glucose declined from 4.83 mmol/L in the first hour after sleep onset to 4.48 mmol/L at 4 to 6 hours (Figure 8B). Over the same hours, the share of deep sleep scored by Ultrahuman fell from 47% in the first two hours to 8% after four hours, while the share of REM sleep rose from 6% to 35% (Figure 8C). Lower glucose during REM sleep therefore coincides with the later part of the night, when glucose is lower anyway. The same applies to Garmin Body Battery, which rises through the night (r=0.69 with hours since sleep onset, excluding the first night) while glucose falls (r=−0.38); this explains its within-sleep correlation with glucose (r=−0.52, Table 3). Ultrahuman skin temperature did not change with hours since sleep onset (r=−0.03), so its within-sleep correlation with glucose (r=0.27) is not explained by the time of night. Total sleep time showed no clear relationship with CV% or TIR on the same date on either device (|r| ≤ 0.23; Figure 6).

**Figure 8.**
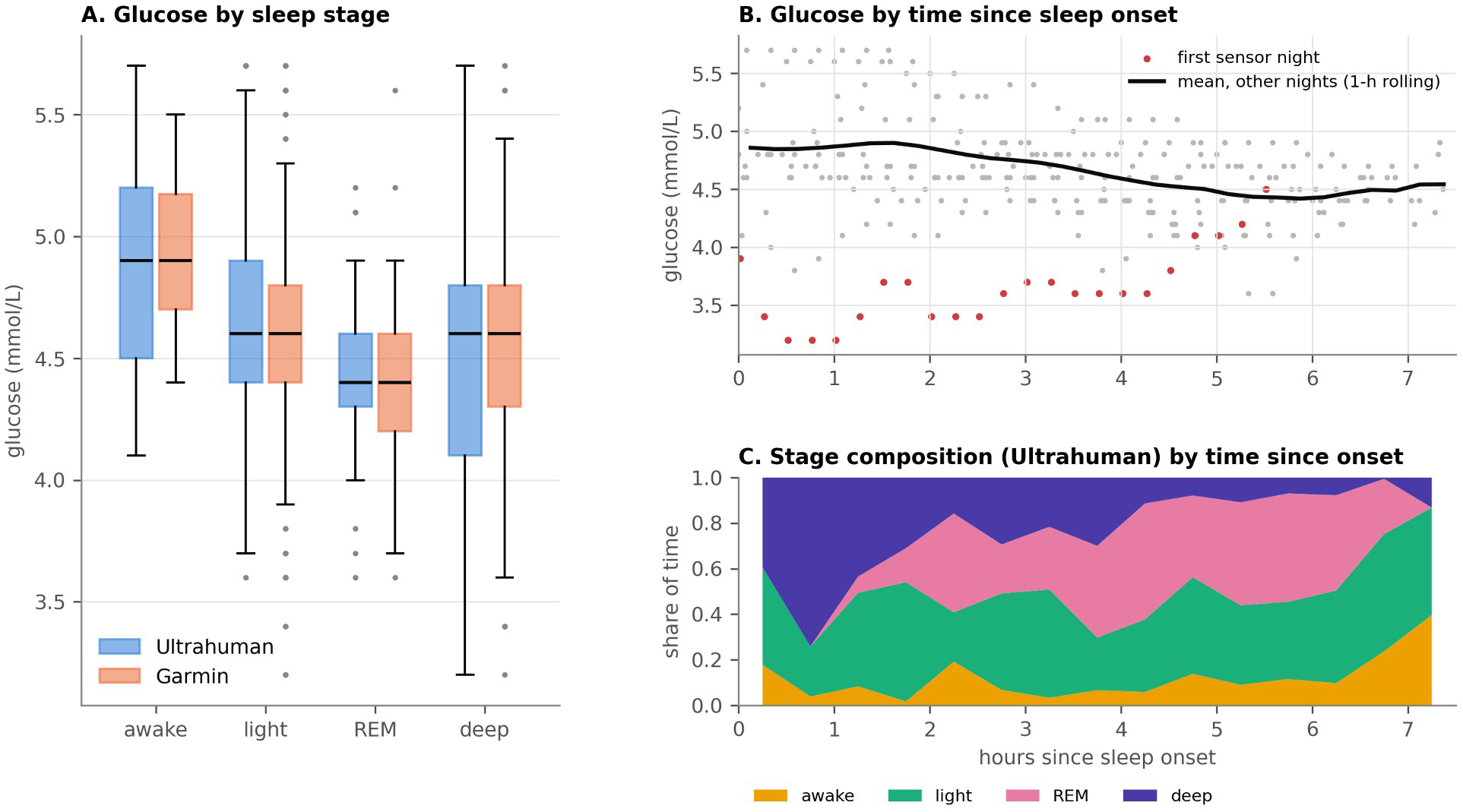
Glucose across the night. (A) Glucose at time points scored as awake, light, REM and deep sleep, by Ultrahuman (blue) and Garmin (orange) staging; boxes show median and interquartile range, whiskers 1.5× the interquartile range. (B) Glucose by hours since Ultrahuman-recorded sleep onset; grey dots, readings from nights after the first; red dots, first sensor night; black line, mean of the other nights in 15-minute bins, smoothed over 1 hour. (C) Share of time in each Ultrahuman sleep stage by hours since sleep onset (30-minute bins, all 12 nights).

## Discussion

### Primary findings

In this n-of-1 study, the Garmin watch and the Ultrahuman ring agreed closely on how long and when the participant slept, but only fairly on sleep stages (kappa 0.28 minute by minute), and not at all on minutes of deep sleep. Their step counts correlated strongly but differed by a large, consistent offset. Heart-rate agreement depended on state and period: moderate during sleep and, before the gap in Ultrahuman data, moderate while awake, but absent while awake after the gap, when the ring’s daytime heart rate stayed near 70 bpm. Heart rate was the wearable signal most closely associated with glucose, but much of that association reflected the day/night rhythm that both share; before the gap the two devices were alike in this respect; and the daily associations rested largely on the first night of sensor wear and on a TIR that varied on only four dates.

### Where the watch and the ring disagreed, and why

Good agreement on sleep duration combined with poor agreement on sleep stages is the pattern seen in studies against PSG, in which consumer wearables detect sleep well but classify its stages less consistently (Chinoy et al., 2021; Kainec et al., 2024; Stone et al., 2020). In the six-device comparison against PSG, kappa ranged from 0.21 for the Garmin Vivosmart 4 to 0.53 for the Apple Watch Series 8, and the Garmin device scored about 42% of PSG-defined REM sleep as light sleep (Schyvens et al., 2025), the same confusion that dominates our Figure 4A. Our kappa between the two devices falls within that range but cannot be compared with it directly: it compares two devices with each other rather than one device with PSG, the Ultrahuman ring belongs to a different class of device, and the Garmin models differ. The hypnograms add that the disagreement was concentrated in the second half of the night, when light and REM sleep alternate. The step offset (about 22% of the two-device mean, with Garmin higher on every day) may reflect wrist versus finger placement and different step-counting algorithms; it is not an artefact of the day the ring was off, which we excluded. Garmin step counts have generally shown good validity against reference measures (Evenson and Spade, 2020), but we found no published validation of Ultrahuman step counts, so the offset cannot be attributed to either device. Part of the 4.3 bpm difference in resting heart rate may come from how each device defines it: Garmin reports the lowest 30-minute average in a 24-hour period (Dial et al., 2025), a minimum rather than an average, whereas Ultrahuman does not document its calculation, so the two indices need not measure the same quantity. Against ECG, Garmin HRV agreed less closely than ring-based HRV (Dial et al., 2025), so the lower Garmin night HRV may partly reflect measurement error on the Garmin side. Heart rate showed a different pattern, as the differences within the day followed sleep/wake state and date. During sleep, the devices differed by a stable few beats per minute. While the participant was awake, and particularly after the Ultrahuman data gap on 3 August, Ultrahuman’s daytime heart rate stayed near 70 bpm regardless of activity, whereas Garmin’s rose with activity. A change in ring fit or measurement mode after the ring was put back on would produce this pattern, but our data cannot separate these explanations from a change in the algorithm, and Ultrahuman’s documentation does not describe how daytime heart rate is sampled or smoothed. The two devices are also worn differently: the watch strap can be loosened or tightened, whereas the ring has a fixed size and cannot be loosened, so the ring’s contact with the skin, and with it the optical signal, may change with finger swelling, temperature and hand position in a way the wearer cannot adjust. A user comparing the two devices during the day would see differences of 20 bpm or more with no indication of which device is right. Daytime heart rate should therefore not be pooled across devices, or across periods of wear of one device, without checking for this kind of shift. In our data the shift also reached the glucose analyses: pooled over the whole period, Ultrahuman’s waking heart rate seemed to carry no glucose signal, but before the gap it matched Garmin’s.

### What the glucose associations reflect

Heart rate was more closely associated with glucose than sleep stages were, but separating sleep from waking hours shows how much of the overall association came from the day/night rhythm. The Garmin heart-rate correlation fell from 0.43 to 0.34 within sleep and 0.21 within waking hours. The two devices did not differ in this respect before the Ultrahuman data gap (waking r=0.25 and 0.24); Ultrahuman’s near-zero waking correlation over the whole period is a consequence of the ring’s behaviour after the gap, not a property of ring-based heart rate. Skin temperature is the clearest case of the day/night confounding, as its correlation with glucose changed sign once sleep and waking hours were analyzed separately. Within sleep, the time of night confounds in the same way: Garmin Body Battery rises through the night while glucose falls, which produces its within-sleep correlation with glucose without any direct link. The direction of the heart-rate association also argues against a response to glucose itself: here lower glucose went with lower heart rate, whereas in insulin-treated diabetes heart rate rises during hypoglycemia (Lehmann et al., 2023). The meal-by-meal analysis showed a larger Garmin heart-rate rise after the meals and coffees that produced a larger glucose rise, and no such pattern for Ultrahuman even before the gap, but most meal windows overlapped another event and the pattern was weak among the events with a clean window, so we regard it as exploratory. Heart-rate peaks at meal onset have been recorded with a Garmin watch in people with post-bariatric hypoglycemia (Cossu et al., 2026), and our data cannot tell whether glucose and heart rate respond to a common driver, such as the size of the meal, or whether one affects the other. Across days, the largest correlations depended on the two dates at the start of sensor wear, when CGM readings are least accurate (Tsoukas et al., 2020) and when pressure on the sensor, for example from lying on it, can produce false overnight lows (Mensh et al., 2013); the correlations with TIR are further limited by a TIR that was 100% on 8 of the 12 dates. Lower glucose in REM sleep is likewise better explained by the time of night than by the stage itself. A study linking Garmin sleep stages to nocturnal hypoglycemia in type 1 diabetes found more deep and REM sleep on nights with hypoglycemia but did not report adjusting for time of night (Alsuhaymi et al., 2026). Taken together, our data do not show that heart-rate, HRV or sleep-stage signals can serve as a device-independent indicator of glucose.

### Strengths and limitations

The main strength of this study is that two consumer wearables with independent, proprietary algorithms and a CGM recorded the same person over the same period in daily life, and that we analyzed the data by sleep/wake state, time of day, date and period of wear as well as in aggregate. The main limitation is the n-of-1, 12-day design: the findings describe one participant’s devices and physiology and cannot be generalized, and no sample size was planned. The daily analysis rests on 12 consecutive dates from one person, and the intraday analyses on autocorrelated series, so the numbers of readings overstate the independent information and we have not tested any correlation for significance; the 39 daily coefficients should be read as a description of this record. With 12 days, the Bland-Altman limits of agreement are imprecise (each limit has a 95% CI of about ±1.1 SD of the differences). Neither wearable was compared with a reference such as PSG or ECG, so our results describe how consistent the devices were with each other, not how accurate they were, and we found no peer-reviewed validation of the Ultrahuman ring against PSG or ECG. The ring’s data gap and changed daytime heart rate after 3 August affected half of the record, and although we report the two periods separately, the cause of the change is unknown. The CGM record covers 253.5 hours, less than 12 full days and below the 14 days recommended for reliable CGM metrics (Battelino et al., 2019), with four gaps of more than one hour. TIR and CV% were computed over all readings, including scans taken whenever the participant chose, which gives scanned periods more weight than a time-weighted calculation would. Finally, the TIR range of 3.9 to 10.0 mmol/L was designed for diabetes and discriminates little in a person whose glucose never exceeded 10.0 mmol/L. For comparison, the participant’s CV of 16.0% was close to the mean of 17% reported in healthy children and adults without diabetes, measured with a different sensor (Shah et al., 2019).

## Conclusions

Two consumer wearables worn together agreed on how long this participant slept but not on how the night was staged. Their heart-rate readings agreed during sleep and, before one device’s data gap, while awake, but diverged during the day after the gap. Heart rate tracked glucose more closely than any sleep metric did, but much of that association came from the day/night rhythm that both share, and the daily associations depended on the first night of sensor wear. As hypotheses for larger studies rather than established guidance, we suggest that studies which combine data from different wearables, or relate wearable data to CGM, analyze sleep and waking hours separately, check device agreement over the whole period of wear, and exclude or model the first day of each CGM sensor. Replication in more people over longer periods, with a validated reference for at least one wearable signal, is needed before these patterns can be generalized.

## Funding

This study received no external funding.

## Conflicts of Interest

The authors declare no conflict of interest.

## Data Availability

The data and analysis code supporting this study are available from the corresponding author on reasonable request.

## Use of Generative AI

Claude (Anthropic) was used for coding support in the data analysis and for writing support during the preparation of this manuscript. The authors reviewed and edited all code and text and take full responsibility for the content of the publication.

## Abbreviations

API: application programming interface
bpm: beats per minute
CGM: continuous glucose monitoring
CI: confidence interval
CV%: coefficient of variation
ECG: electrocardiography
HR: heart rate
HRV: heart rate variability
IQR: interquartile range
LoA: limits of agreement
LODO: leave-one-day-out
PSG: polysomnography
REM: rapid eye movement
RQ: research question
SD: standard deviation
TIR: time in range
UH: Ultrahuman
VO2max: maximal oxygen uptake

## Notes

### Competing Interest Statement

The authors have declared no competing interest.

